# *cellSight:* Characterizing dynamics of cells using single-cell RNA-sequencing

**DOI:** 10.1101/2025.05.16.654572

**Authors:** Ranojoy Chatterjee, Chiraag Gohel, Ali Reza Taheriyoun, Brett A Shook, Ali Rahnavard

**Affiliations:** Computational Biology Institute, Department of Biostatistics and Bioinformatics, Milken Institute School of Public Health, The George Washington University, Washington, DC 20052; Department of Biochemistry and Molecular Medicine, School of Medicine and Health Sciences, The George Washington University, Washington, DC 20052

**Keywords:** single-cell analysis, intercellular communication, cell interaction

## Abstract

Single-cell analysis has transformed our understanding of cellular diversity, offering insights into complex biological systems. Yet, manual data processing in single-cell studies poses challenges, including inefficiency, human error, and limited scalability. To address these issues, we propose the automated workflow *cellSight*, which integrates high-throughput sequencing in a user-friendly platform. By automating tasks like cell type clustering, feature extraction, and data normalization, *cellSight* reduces researcher workload, promoting focus on data interpretation and hypothesis generation. Its standardized analysis pipelines and quality control metrics enhance reproducibility, enabling collaboration across studies. Moreover, *cellSight*’s adaptability supports integration with emerging technologies, keeping pace with advancements in single-cell genomics. *cellSight* accelerates discoveries in single-cell biology, driving impactful insights and clinical translation. It is available with documentation and tutorials at https://github.com/omicsEye/cellSight.

## INTRODUCTION

The advent of high-throughput single-cell technologies has generated unprecedented volumes of biological data, presenting significant computational challenges in processing, analyzing, and interpreting cellular heterogeneity and dynamics^1–4^. The field of single-cell genomics emphasizes the growing need for sophisticated computational tools^5,6^ capable of handling the increasing complexity of data analysis, particularly in understanding cellular heterogeneity^7–9^ and their temporal dynamics. The computational demands are further amplified by recent technological advances in single-cell methodologies, such as droplet-based^10,11^ and plate-based assays^12,13^, which can generate data from thousands to millions of individual cells in a single experiment.

The computational complexity of single-cell analysis extends beyond mere data volume^14,15^. Current analytical pipelines must address multiple challenges, including batch effect correction, dimensionality reduction, feature selection, and cell type identification^16,17^.

Traditional computational approaches often rely on manual quality control (QC) steps and differential expression analyses, which are computationally intensive and prone to reproducibility issues and human error^18–20^. These challenges are compounded by the high-dimensional nature of single-cell data, where each cell is characterized by the expression levels of tens of thousands of genes^21,22^.

To address these computational challenges, we introduce *cellSight*, an innovative automated computational framework specifically designed to handle the complexity and scale of single-cell data analysis^23,24^. Our solution integrates state-of-the-art computational methods for data processing, visualization, and interpretation. Central to *cellSight* architecture is its robust QC pipeline integrated with statistical modeling and differential expression analyses by considering the usual linear and generalized linear models, emphasizing the role of zero-inflation in the interval data. The results are used in the pipeline to analyze intercellular communication networks during homeostasis.

*cellSight* represents a significant advance in computational biology, providing an automated, end-to-end solution that addresses key computational bottlenecks in single-cell analysis^25^.

The framework incorporates parallel processing capabilities and optimized algorithms for handling large-scale datasets while maintaining statistical rigor and biological relevance. By automating complex computational tasks, *cellSight* not only enhances the efficiency of single-cell analysis but also improves reproducibility and reduces computational overhead^26^.

To validate our computational framework, we applied *cellSight* to two distinct datasets: a mouse skin injury model and a previously published skin aging study^27^. Our automated pipeline successfully reproduced the findings from the skin aging study while generating novel insights into the role of fibroblasts during the healing process in the mouse injury model. These results demonstrate the robustness and versatility of our computational approach in handling diverse biological contexts while maintaining computational efficiency.

Building upon traditional single-cell RNA sequencing analysis, *cellSight* incorporates advanced spatial transcriptomics capabilities through a novel Graph Attention Convolution (GATconv) module. This integration addresses the growing need to understand not only cellular heterogeneity but also the spatial organization and neighborhood effects that drive tissue function and pathology^28,29^.

The spatial module leverages graph neural network architectures to analyze spatially-resolved transcriptomic data^30,31^, where each cell or spot is represented as a node with associated gene expression profiles and spatial coordinates. The methodology employs self-attention mechanisms^32,33^ to weight the importance of neighboring cells based on both transcriptional similarity and spatial proximity. This dual consideration enables the identification of spatially-constrained gene expression patterns and cellular communication networks that would be missed by conventional single-cell analysis approaches^34,35^.

The GATconv implementation in *cellSight* constructs adjacency matrices based on spatial relationships^36^, allowing the model to learn both local neighborhood effects and broader spatial patterns across tissue architecture. Through iterative attention mechanisms^37,38^, the module identifies spatial domains, tissue boundaries, and gradient expression patterns that reflect underlying biological organization. This capability is particularly valuable for understanding developmental trajectories^39^, disease progression patterns^40,41^, and the spatial context of cellular interactions within complex tissues^42,43^.

By integrating spatial transcriptomics with *cellSight*’s existing single-cell analysis pipeline, researchers can seamlessly transition between cell-centric and spatially-aware analyses within a unified computational framework^44,45^. This integration enables comprehensive characterization of tissue organization, from individual cell states to multicellular spatial domains, providing unprecedented insights into the relationship between cellular identity and tissue architecture^46,47^.

## RESULTS

### DYNAMICS OF CELLS IN INJURY-INDUCED SKIN INFLAMMATION

An epithelium covers the outer layer of skin on all animals, protecting against damage, water loss, and infection. When the epithelial layer is damaged, the body swiftly induces a plethora of molecular and cellular changes to repair the tissue. The process of healing has been characterized as three major overlapping phases^48–50^. The first stage is inflammation, which is prompted as the tissue is damaged, involves the recruitment and activation of pro-inflammatory immune cells. The second phase involves the creation of granular tissue and a new epithelium over the injured area. The final stage is the remodeling of the newly generated tissue, particularly the extracellular matrix. The healing process is a time-intensive process that can take months.

Four distinct datasets were meticulously collected from a comprehensive study involving two groups of mice: two datasets from injured skin and two from uninjured skin. This meticulous selection of datasets aimed to unravel the intricate molecular roles and dynamics occurring during the process of injury. Each dataset represents a snapshot of the cellular landscape at a specific point in time, capturing the unique gene expression profiles and cellular responses associated with injury and contrasting them with those from uninjured samples. Including injured and uninjured samples provides a valuable comparative framework, allowing researchers to pinpoint differentially expressed genes and pathways specific to the injury context. This approach enables a thorough exploration of the molecular mechanisms underlying the response to injury, potentially revealing critical insights into the cellular processes, signaling pathways, and dynamic changes in gene expression that contribute to tissue repair or exacerbate damage. The carefully curated datasets form the foundation for a detailed and nuanced analysis, fostering a deeper understanding of the roles played by specific genes and cellular components in the context of injury and recovery in the mouse model.

*cellSight’s* pipeline streamlined the complex single-cell data analysis into a straightforward and efficient process, as demonstrated in our study of skin injury response. Through automated quality control, normalization, and clustering, we successfully detected differentially expressed genes between injured and naive samples, revealing the direct influence of fibroblast subtypes on the wound healing process. The integration module enabled an overarching visualization of the differences in biological states across sample types, while cell-cell interaction analysis systematically mapped the multifaceted communication networks governing the coordinated healing response. This case study demonstrates the ability of researchers to rapidly extract biologically meaningful insights from complex single-cell data without requiring exhaustive computational expertise.

To illustrate the end-to-end automation functionalities of *cellSight*, we ran our developed pipeline on single-cell datasets from a mouse skin injury model. The automated pipeline successfully processed four datasets consisting of 39,466 cells over 32,285 genes, thereby removing the need for manual data manipulation and format conversion, which typically consumes much time for researchers. *cellSight*’sQC module automatically applied optimized filtering thresholds (RNA count >200, unique genes >2500, total RNA molecules >8000, mitochondria percentage = 0) without the need for manual parameter tuning or iterative tuning cycles. The pipeline then successfully performed canonical correlation analysis for dataset integration, a computationally demanding step that typically necessitates expert expertise, and carried out unsupervised clustering that revealed 21 cell populations.

Moreover, *cellSight* automates the creation of publication-ready visual representations in the form of violin plots, feature plots, and dimensional reduction embeddings, thereby greatly speeding up the process of cell type annotation. Although the pipeline does not automatically apply cell type labels, it produces standardized expression matrices and interactive visualizations that enhance the efficiency of expert annotation. Through these automated visualizations, we observed that fibroblast populations expressed *Pdgfr*α, Col1a1, and *Dcn*^*51–53*^, while immune populations were demarcated by markers including *Adgre1* (macrophages) and *Ly6c2* (monocytes) cell types^54–56^. This automation-assisted strategy retains the important role of expert biological knowledge in the identification of cell types while reducing annotation times from days to hours. **Fig. 2A** illustrates how *cellSight* automatically renders extensive visualizations of quality control metrics without manual formatting, while **Fig. 2B** depicts successful multi-sample integration through dimensional reduction visualization, an effort that would otherwise entail extensive coding proficiency.

**Figure 1.**
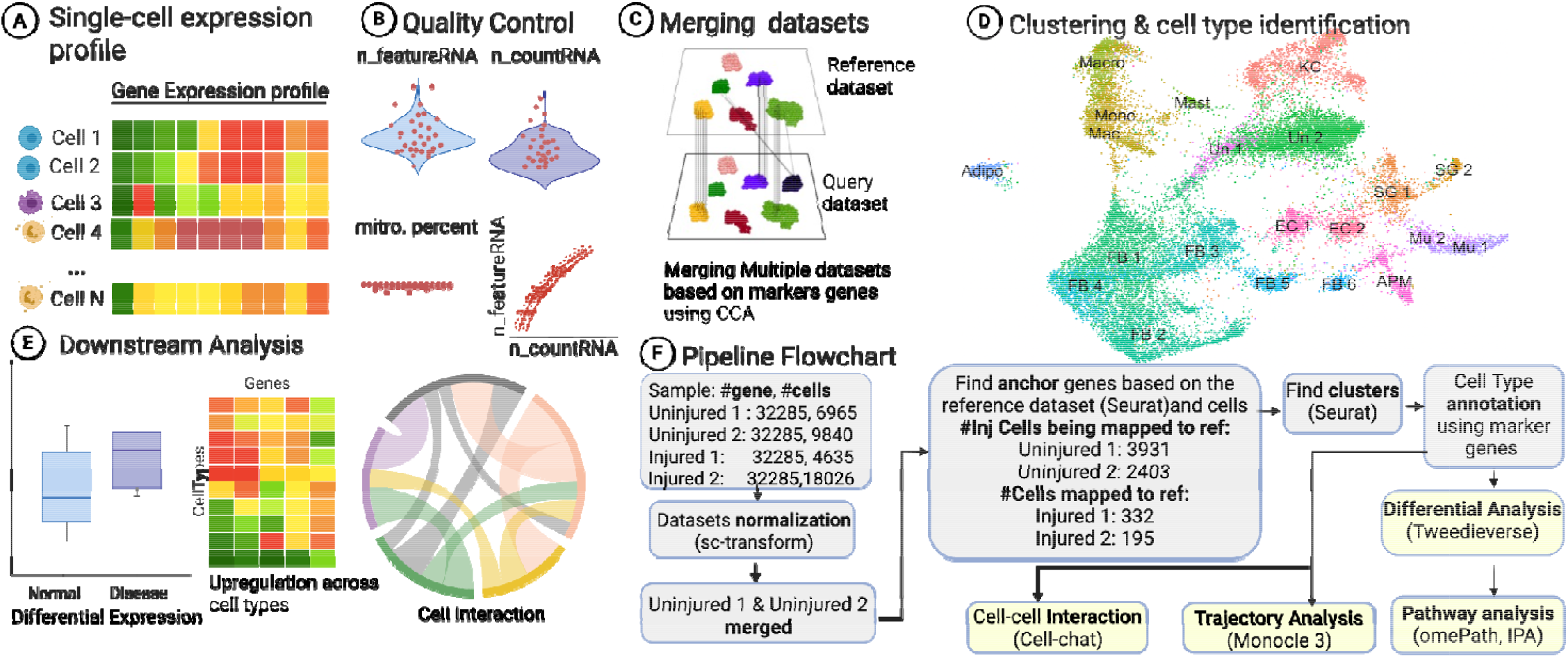
Single-cell Analysis Pipeline for Transcriptional Profiling. (**A**) Single-nucleus RNA expression heatmap displaying gene expression patterns across individual cells (Cell 1, Cell 2,…, Cell N). (**B**) Quality control is based on principal components to distinguish viable cells from unreliable data points. (**C**) Dataset integration through canonical correlation analysis (CCA), merging reference and query datasets to identify common cell populations. (**D**) High-dimensional clustering visualization showing distinct cell type populations with annotated labels (e.g., Endothelial Cells, Fibroblasts) created in the previous step. (**E**) Downstream analyses featuring circular segmented plots of cell-cell interactions and the significance of the disease states visualized by box plots and heatmaps. (**F)** cellSight workflow: Data normalization and integration, cluster annotation, differential expression (Tweedieverse), trajectory analysis (Monocle3), pathway analysis (omePath), and cell-cell interaction mapping (CellChat).

**Figure 2.**
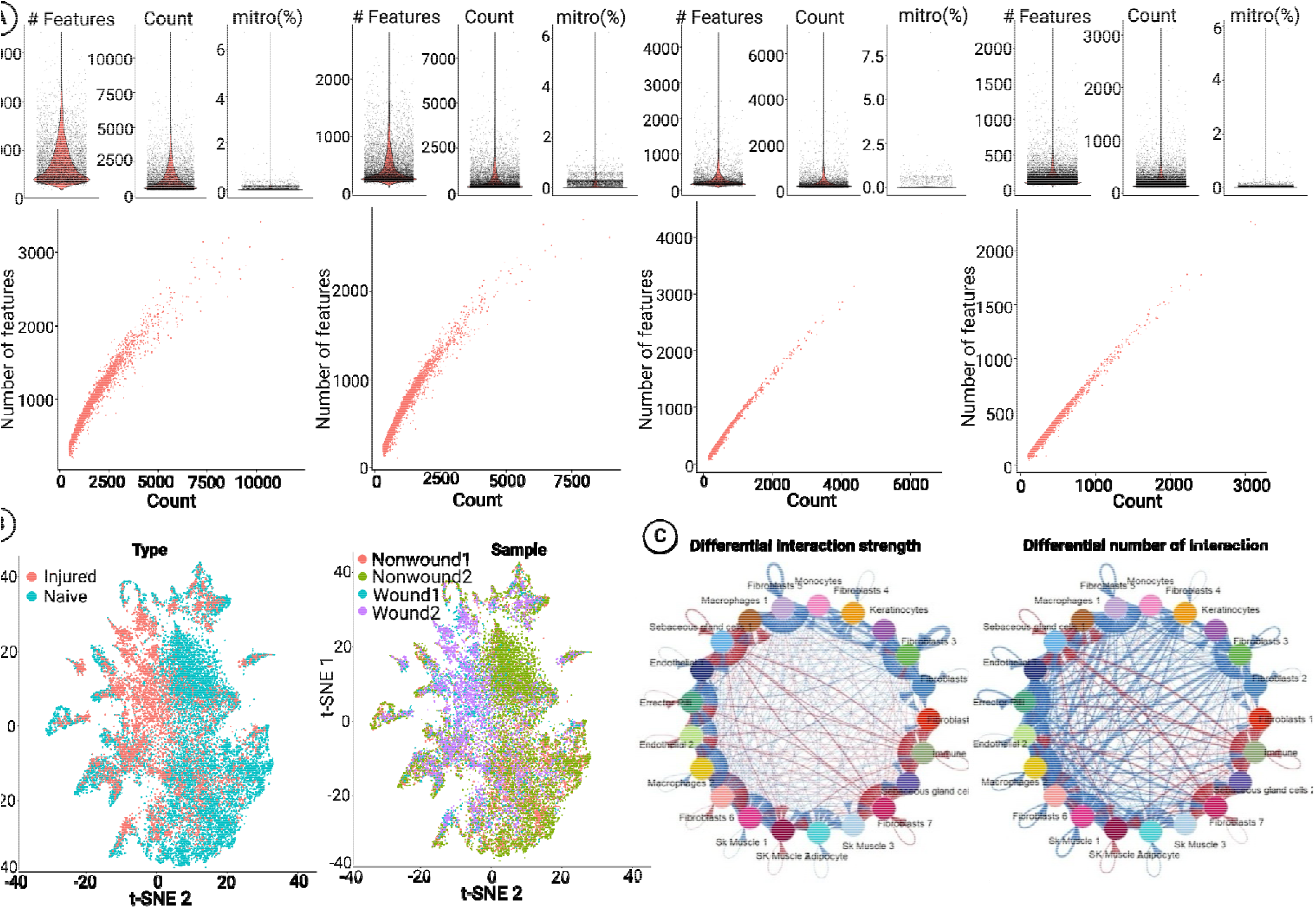
Comprehensive computational analysis of injury-induced transcriptional changes in mouse skin tissue. (**A**) QC metrics assessment, including gene detection rates, RNA molecule quantification, and mitochondrial content analysis. (**B**) Multi-sample integration validation through dimension reduction visualization. (**C**) Cell-cell interaction network analysis highlights the activated communication pathways between keratinocytes, immune cells, and fibroblasts during wound response.

Subsequently, *cellSight*’s differential expression module automatically performs statistically rigorous comparisons between experimental conditions using the Tweedie model, specifically designed to handle the zero-inflation properties inherent in single-cell data. By automating such intricate analytical processes from preprocessing and normalization to visualization and statistical evaluation, *cellSight* removes many of the error-prone, time-consuming manual steps commonly involved in single-cell workflows. This end-to-end automation makes cutting-edge transcriptomic analysis accessible to researchers lacking specialized computational training, thereby democratizing access to the insights of single-cell biology within the biomedical research community.

After the naming of each cluster was completed, differential expression analysis was performed on the 4 samples to find out the genes that have changed in their expression after injury using the Tweedie model. We found several cytokines and growth factors expressed by keratinocytes and monocytes alongside the fibroblast clusters. In addition, we also found some cytokines and growth factors that were uniquely expressed by the fibroblast clusters only. The results clearly show the importance of *Ccl* in injury since *Ccl* (*Ccl2*) is employed to recruit monocytes and macrophages to the wounded area, facilitating healing. Cell-cell interaction network analysis (**Fig. 2C**) identified activated signaling pathways between keratinocytes, immune cells, and fibroblasts during wound response. The results highlight the potential importance of Ccl in injury, with our analysis showing elevated Ccl2 expression, which is known to be involved in recruiting monocytes and macrophages to wounded areas^57,58^. Cell-cell interaction network analysis (**Fig. 2C**) predicted activated signaling pathways between keratinocytes, immune cells, and fibroblasts during wound response. Our analysis suggests that increased *Ccl2* expression could facilitate monocyte/macrophage recruitment^59^, while pro-inflammatory cytokines (*IL-1, IL-6, IL-8*) may initiate immune response and pathogen clearance^60,61^. Based on known pathway functions, the observed *IL-4/IL-13* expression patterns are predicted to promote fibroblast proliferation and collagen synthesis^62^, with IL-8 potentially stimulating endothelial cell migration and angiogenesis^63^.

### ROLE OF FIBROBLASTS IN SKIN AGING

Historically, knowledge about skin cell components has been largely derived from mouse studies, utilizing reporter constructs, lineage tracing, fluorescence-activated cell sorting (FACS), and immunohistochemistry (IHC). These studies identified distinct fibroblast subtypes in different dermal layers, suggesting functional diversity. However, to bolster our understanding of the mechanism in skin tissue, scRNA-seq data were utilized to capture the underlying workings.

In this study^27^, the researchers conducted a comprehensive analysis of human dermal fibroblasts at the single-cell level, focusing on a defined, sun-protected area from both young (25 and 27 years old) and old (53–70 years old) male Caucasian donors. By analyzing over 15,000 cells, the researchers identified four main fibroblast subpopulations with distinct spatial localizations and characteristic functional annotations.

*cellSight* automated QC module generates comprehensive multi-parameter metrics visualizations (**Fig. 3A**) with validated data quality across young (25-27 years) and old (53-70 years) male donors without manual parameter tuning. The integration analysis module of the pipeline performs automated batch effect correction and provides dimension reduction plots showing effective data harmonization (**Fig. 3B**). With its adaptive clustering algorithms, *cellSight* computes unsupervised clustering visualizations at multiple resolutions (**Fig. 3C**), thereby allowing researchers to effortlessly discern discrete cell populations across all donors without necessitating sophisticated computational skills. The automatically generated high-quality visualizations illustrate how *cellSight* accelerates the analytical process, allowing researchers to devote more time to biological interpretations and less to computational ones.

**Figure 3.**
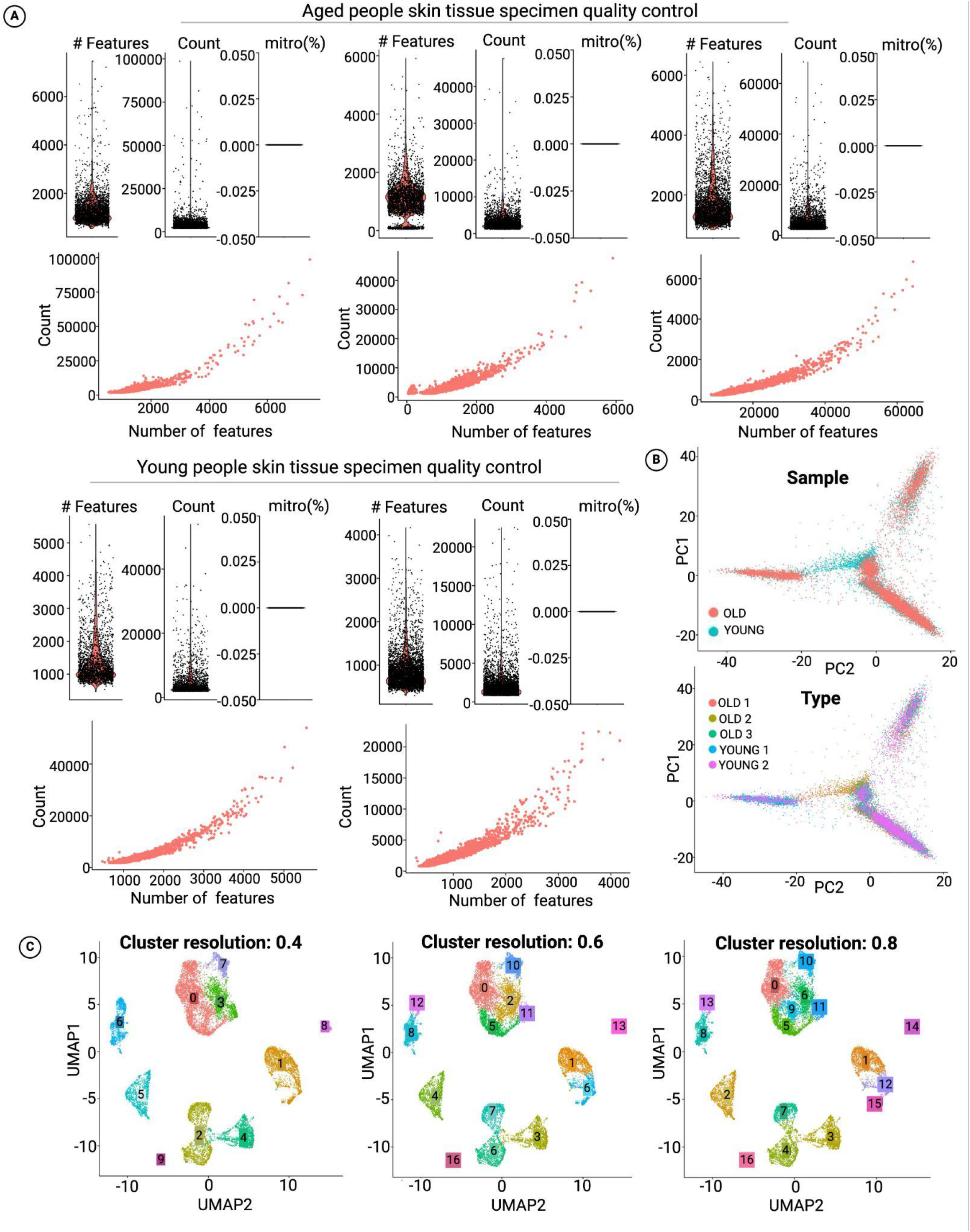
Single-cell transcriptomic analysis of age-dependent changes in human skin fibroblasts. (**A**) Multi-parameter QC metrics demonstrating data quality across samples. (**B**) Sample integration analysis showing batch effect correction and data harmonization. (**C**) Unsupervised clustering analysis at varying resolutions reveals distinct cell populations.

The study provided insights into the differential ‘priming’ of fibroblasts, resulting in functionally heterogeneous subgroups. *cellSight’s* built-in visualization modules automatically produced high-resolution UMAPs, feature plots, and violin plots that reveal this cellular heterogeneity, allowing researchers to identify the age-associated loss of identity among fibroblast subpopulations without any manual parameter optimization. The pipeline also creates standardized differential expression heatmaps that graphically display the decrease in unique features of every subgroup, and its cell-cell interaction visualization tools can generate network diagrams automatically displaying how aged fibroblast subpopulations, which express specific skin aging-associated secreted proteins (SAASP), have predicted decreases in interactions with other skin cell types. The figures present how *cellSight* converts intricate single-cell datasets into easy-to-interpret visualizations that address fundamental transcriptomic inquiries regarding cellular identity, state transitions, and cell-to-cell interactions achieved through an automated pipeline without requiring advanced computational expertise or the creation of custom scripts.

In our second case study, we wanted to verify the proper working of the *cellSight* pipeline. To validate the proper working, we ran the automated pipeline on a previously published study that sampled old and young skin tissue to observe the change in fibroblast priming in old people. Our automated pipeline shows the same results as they obtained from their analysis. In the initial analysis, we integrated cells from all five samples, resulting in a UMAP plot displaying 17 clusters with distinct expression profiles. Notably, each cluster contained cells from all donors. Expression profiling of canonical markers revealed molecular signatures of inflammatory, secretory, and mesenchymal states across fibroblast subpopulations (**Fig. 3C**). Initial analysis integrated cells from all five samples, yielding 17 clusters with distinct expression profiles. Each cluster contained cells from all donors, with fibroblasts, marked by *LUM, DCN, VIM, PDGFRA*, and *COL1A2*^*64–66*^, constituted the most abundant skin cell type, represented by four clusters. Analyzing each sample individually produced a similar number of clusters and identified the same major cell types(**Fig. 4A**).

**Figure 4.**
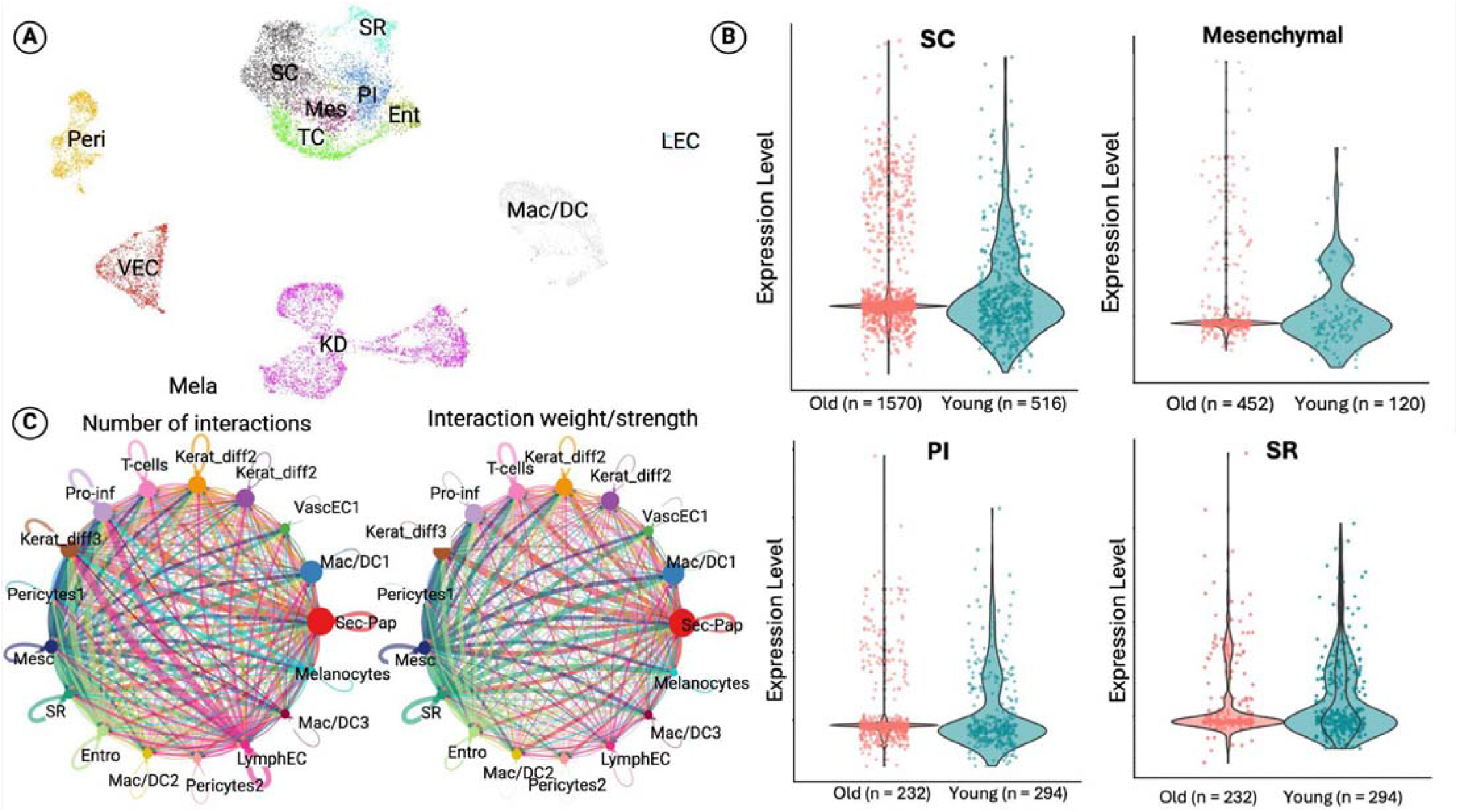
Expression analysis of human skin fibroblasts during aging. (**A**) Cell type identification revealed diverse populations in the skin tissue, including multiple fibroblast subtypes, epithelial cells, immune cells, and vascular components. (**B**) Expression profiling of canonical markers across identified fibroblast subpopulations demonstrates molecular signatures of inflammatory, secretory, and mesenchymal states. (**C**) Intercellular communication network analysis reveals age-associated changes in cell-type specific signaling patterns.

Intercellular communication network analysis (**Fig. 4C**) revealed age-associated changes in cell-type-specific signaling patterns. Key findings included differential *‘priming’* of fibroblasts into functionally distinct subgroups and age-related loss of identity across all fibroblast subpopulations (**Fig. 4B**). Notably, older fibroblast subpopulations expressed specific (SAASP) and showed decreased interactions with other skin cell types.Individual sample analysis produced consistent clustering patterns and cell type identification, validating the robustness of our analytical approach. This comprehensive characterization provides insights into age-related changes in fibroblast function and intercellular communication networks.

As shown in the step-by-step timeline (**Fig. 5E**), *cellSight* condenses ten years of analytic software development (2015-2024) that integrates approaches from early platforms like *Seurat* to the latest methods in spatial transcriptomics and intercellular communication. The end-to-end pipeline addresses the main bottlenecks in single-cell analysis through automated quality control, normalization, dimensionality reduction, clustering, and biological interpretation. *cellSight* is unique in that it integrates *Tweedieverse*^*67*^ for differential expression analysis and *CellChat* for intercellular communication networks, benchmarked with fibroblast heterogeneity characterization in wound healing and aging models. The pipeline overcomes key analytical hurdles such as batch effect correction, parameter optimization, and cell type annotation without compromising statistical stringency and biological relevance. By reducing computational skill demands at the cost of analysis trade-offs, *cellSight* provides workflow standardization for use across varied tissue sources and experimental setups. With a scalable design tuned for managing datasets ranging from thousands to millions of cells, *cellSight* facilitates reproducibility and enables advanced single-cell analytics for the wider research community.

**Figure 5.**
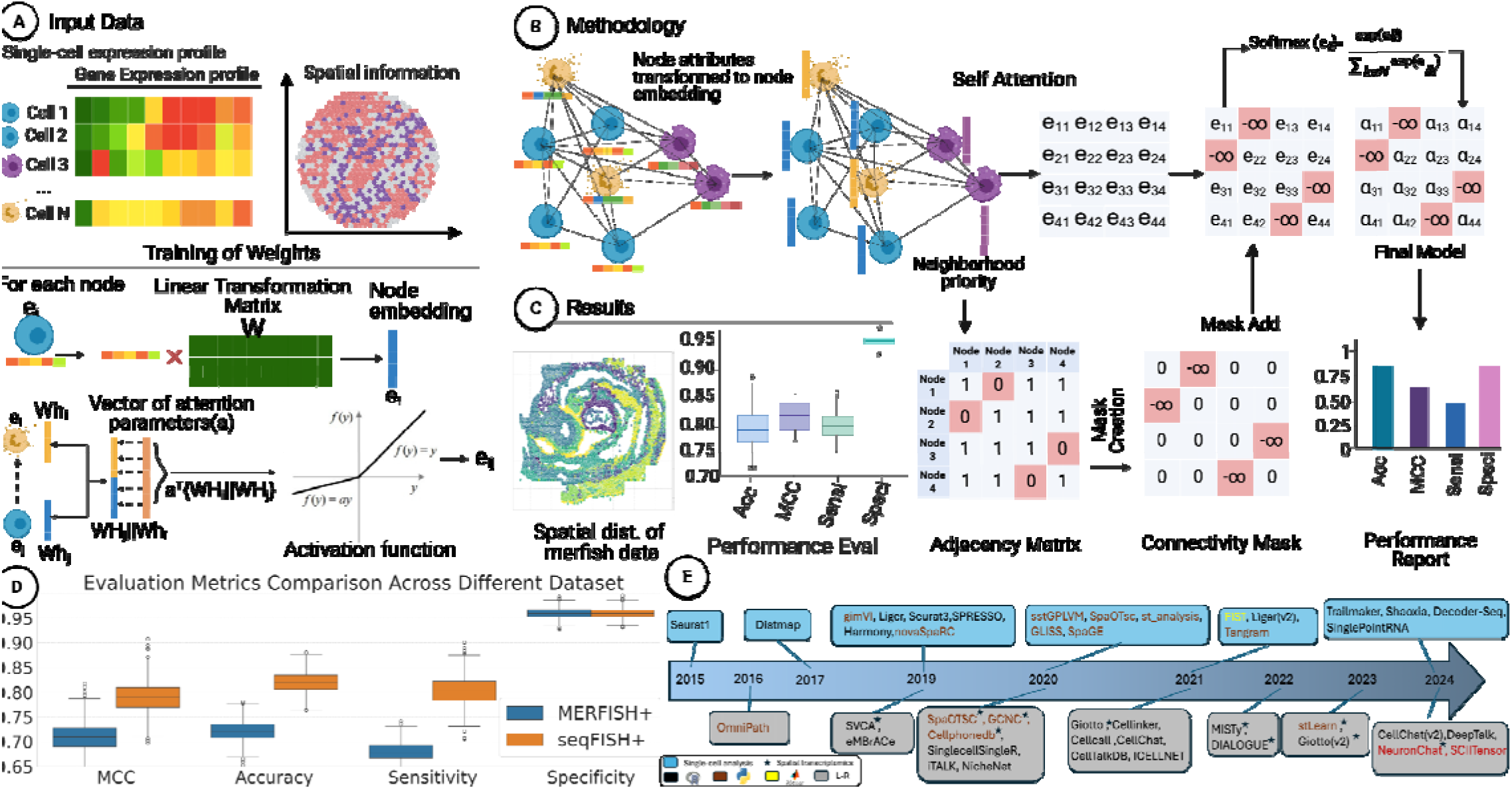
Computational framework for spatially-aware graph neural network analysis of single-cell expression data and timeline of analytical tool development (2015-2024). (**A**) Input data comprises single-cell gene expression profiles and spatial coordinate information from tissue sections. (**B**) Methodology illustrates the graph neural network architecture where node attributes are transformed into embeddings through linear transformation. The self-attention mechanism computes attention weights between connected nodes using softmax, with mask addition operations (indicated by values) controlling information flow through the network. (**C**) Results show spatial visualization of merfish data and performance metrics including accuracy (Acc), Matthews correlation coefficient (MCC), sensitivity (Sensi), and specificity (Speci). The adjacency matrix and connectivity mask illustrate spatial relationships between nodes. (**D**) Quantitative comparison between *MERFISH+* (blue) and *seqFISH++* (orange) datasets demonstrating consistent performance across platforms. (**E**) Timeline spanning 2015-2024 showing evolution from foundational platforms (Seurat1, Distmap) through specialized spatial methods (*gimVI, Liger, SeuratV3*) to advanced integration tools (*stGPLVM, SpaOTsc, Tangram, CellChat, Giotto, stLearn, SCITensor*). Asterisks indicate spatial transcriptomics-specific methods, illustrating the field’s progression toward integrated, spatially-aware analytical frameworks.

### LIGAND-RECEPTOR INTERACTION USING GATCONV

To validate the effectiveness of our Graph Attention Convolution (GATconv) module for spatial transcriptomics analysis, we conducted comprehensive benchmarking against established graph-based approaches and spatial transcriptomics methods. We evaluated GATconv’s performance against graph convolutional neural networks for genes (GCNG) using four key metrics: Matthews correlation coefficient (MCC), accuracy, sensitivity, and specificity across multiple independent runs (**Fig. 5C**).

GATconv demonstrated superior performance across all evaluation metrics compared to GCNG. The method achieved higher accuracy with a median value of 0.85 (IQR: 0.84-0.86) compared to GCNG’s median of 0.80 (IQR: 0.79-0.81), representing a 6.25% improvement (**Fig. 5C**). The MCC showed GATconv performing at 0.78 (IQR: 0.77-0.79) versus GCNG at 0.75 (IQR: 0.74-0.76). For sensitivity, GATconv reached 0.82 (IQR: 0.81-0.83) while GCNG achieved 0.80 (IQR: 0.79-0.81), indicating enhanced ability to detect true positive interactions. Both methods maintained comparable specificity at approximately 0.97.

We further assessed GATconv’s computational stability across different scales by testing performance at 1,000 runs, 100,000 runs (variant 1), and 100,000 runs (variant 2). The analysis revealed consistent performance across all computational scales. At 1,000 runs, GATconv maintained MCC values with a median of 0.75 (IQR: 0.74-0.76) and accuracy of 0.80 (IQR: 0.78-0.82). When scaled to 100,000 runs, both variants demonstrated nearly identical performance distributions with MCC values clustering around 0.72-0.73 and accuracy maintaining 0.82-0.83 ranges. This stability indicates robust algorithmic performance suitable for large-scale spatial transcriptomics applications.

Additionally, we evaluated GATconv’s performance against established spatial transcriptomics methods, specifically *MERFISH+* and *seqFISH+* (**Fig. 5D**). The comparative analysis assessed the module’s ability to process spatially-resolved gene expression data while maintaining computational efficiency in ligand-receptor interaction prediction. The benchmarking results demonstrated that GATconv maintained competitive performance when compared to these specialized spatial transcriptomics approaches, showing consistent accuracy across different spatial resolution scales and indicating adaptability to various spatial transcriptomics platforms and data structures (**Fig. 5A-B, D**).

## DISCUSSION

The *cellSight* platform is an important development in single-cell RNA sequencing analysis that provides investigators with an extensible and automated pipeline incorporating a wide range of analytical methods into one platform, as demonstrated in **Fig. 6**. The pipeline initiates with stringent quality control and preprocessing procedures prior to creating insightful violin plots (**Fig. 6A)** that reveal the distribution of gene expression in the detected clusters. These meticulously crafted plots simultaneously depict expression prevalence and intensity, where the width precisely captures the proportion of cells expressing particular markers at specific levels while the height corresponds to expression magnitude. This dual visualization enables researchers to immediately discern cluster-specific expression patterns across multiple cell types, revealing both subtle and pronounced differences that define cellular identities without requiring labor-intensive manual annotation procedures.

**Figure 6.**
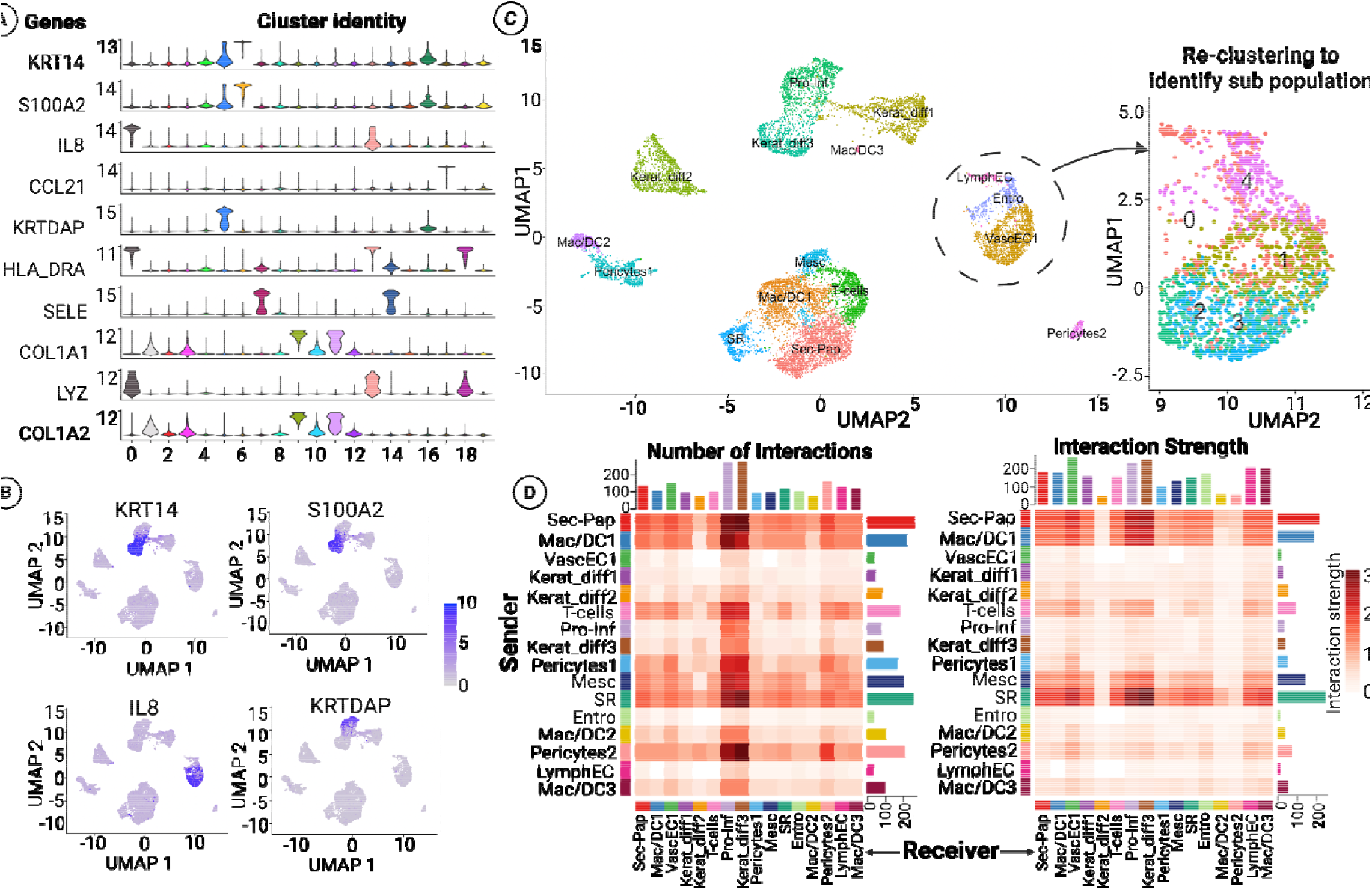
Multi-dimensional analysis of cellular heterogeneity and intercellular interactions within tissue microenvironment. (**A**) Violin plots showing marker gene expression distributions across cell clusters. (**B**) Feature plots mapping marker expression (blue intensity) on UMAP coordinates. (**C**) UMAP visualizations of cell clustering - left panel shows major cell populations, right panel shows five subclusters (0-4) within a specific region. This hierarchical approach is essential for identifying rare cell states, differentiation intermediates, and disease-associated variants that play critical functional roles despite representing small fractions of the tissue. (**D**) Cell-cell interaction analysis through paired heatmaps quantifying communication frequency (left) and strength (right) between sender (y-axis) and receiver (x-axis) populations. These matrices reveal the intercellular signaling networks orchestrating tissue function, highlighting key communication channels and regulatory hubs that coordinate multicellular responses in development, homeostasis, and disease.

Complementing this approach, feature plots (**Fig. 6B**) project expression patterns onto two-dimensional UMAP coordinates with remarkable precision. The gradient-based color intensity systematically highlights expression hotspots across the cellular landscape, creating intuitive spatial maps that reveal both localized expression domains and transitional zones between cell states. These visualizations transform complex multidimensional expression data into interpretable spatial relationships that facilitate the identification of functional territories and developmental trajectories within heterogeneous tissues.

The UMAP visualization module (**Fig. 6C**) demonstrates *cellSight’s* sophisticated dimensionality reduction and clustering algorithms by depicting how transcriptionally similar cells aggregate within high-dimensional space. The left panel presents a comprehensive overview of the cellular ecosystem with precisely labeled populations reflecting their distinctive transcriptional signatures. The right panel showcases the pipeline’s re-clustering functionality, which addresses a critical challenge in single-cell analysis, the detection of rare or transitional cell states that often remain hidden within broader populations. This re-clustering approach enables targeted analysis of specific cellular compartments (illustrated by the circled vascular region) to reveal cryptic subpopulations with distinct molecular signatures. Such hierarchical subpopulation discovery is particularly valuable for identifying stem cell niches, differentiation intermediates, and disease-associated cell states that might represent only a fraction of the broader population yet play pivotal roles in tissue function.

The ability to iteratively re-cluster specific regions of interest allows researchers to progressively refine cellular taxonomies, uncovering biological heterogeneity at multiple resolutions that more accurately reflect the complex organizational hierarchies present in living tissues. This capability proves especially valuable in disease contexts where pathological cell states may emerge as subtle variants of normal populations, enabling more precise identification of therapeutic targets and biomarkers.

A particularly innovative component of *cellSight* is the integrated module for cell-cell interaction analysis (**Fig. 6D)**, generating paired heatmaps that quantitatively assess both the frequency and strength of predicted molecular communications between distinct cellular populations. These comprehensive interaction matrices, with sender populations arrayed vertically and receiver populations horizontally, systematically capture the elaborate intercellular signaling networks that orchestrate tissue function. The complementary metrics of interaction frequency and strength provide multidimensional insights into communication dynamics, highlighting predominant signaling axes and revealing potential regulatory mechanisms underlying tissue homeostasis or disease progression. This systematic approach to mapping cellular crosstalk uncovers critical signaling nodes and potential therapeutic targets that would remain undetected through conventional gene expression analysis alone.

In the mouse injury study, the analysis of wounded and naive skin tissue samples revealed significant insights into the role of fibroblasts and other cell types in wound healing. Our discovery of seven distinct fibroblast subtypes (FB1-FB7) emphasizes the functional heterogeneity within this population. This suggests that fibroblasts play diverse roles in different stages of tissue repair, potentially contributing to processes such as extracellular matrix remodeling^68^, cytokine secretion^69,70^, and cellular signaling^71^ during healing.

Detecting differentially expressed genes between the injured and naive samples, particularly those encoding cytokines and growth factors, further supports the pivotal role of fibroblasts and other immune cells (monocytes and macrophages) in modulating the inflammatory response. The upregulation of *Ccl2* is of particular interest as it reinforces the importance of chemokine-mediated recruitment of immune cells to the wound site. Similarly, the observed increase in pro-inflammatory interleukins (*IL-1, IL-6, IL-8*)^72^ aligns with their well-established roles in initiating the early phases of inflammation, immune cell recruitment, and tissue repair. These findings align with current knowledge of wound healing, where an orchestrated sequence of cellular communication is essential for efficient repair. Identifying specific fibroblast-expressed cytokines^73^ and growth factors^74,75^ unique to this cell population opens new avenues for understanding how fibroblasts regulate their local environment and contribute to effective tissue repair.

In the skin aging study, the focus shifted to understanding how fibroblast identity and functionality change with age. By validating the cellSight pipeline on a previously published dataset, we confirmed the age-related decline in the distinct characteristics of fibroblast subtypes. Notably, older fibroblast populations exhibited reduced functional specialization, coupled with the expression of aging-associated secreted proteins (SAASP)^76,77^, consistent with the senescence-related changes in cellular communication observed in aged tissues^78^. The loss of fibroblast identity observed in aged donors is concerning as it suggests that fibroblasts lose their ability to efficiently participate in tissue homeostasis and repair over time, potentially contributing to age-related tissue dysfunction.

Our findings are consistent with previous studies that have linked fibroblast dysfunction with impaired wound healing and tissue regeneration in older individuals. The decreased interactions between fibroblasts and other skin cell types in aged samples, as predicted by the cell-cell communication analysis, may explain the diminished regenerative capacity of aging skin. Moreover, the presence of SAASP in old fibroblasts likely contributes to the chronic low-grade inflammation associated with skin aging, further impairing tissue repair mechanisms.

Taken together, these two studies underscore the critical roles that fibroblasts play in both wound healing and skin aging. In the context of injury, fibroblasts facilitate repair by promoting immune cell recruitment, extracellular matrix deposition, and tissue remodeling. However, during aging, fibroblasts progressively lose their functional identity, contributing to impaired healing and increased tissue fragility. These insights have important implications for developing therapeutic strategies aimed at enhancing wound healing and mitigating the effects of aging on skin repair.

Alongside biological insights, *cellSight’s* modular architecture is expressly configured to facilitate the incorporation of novel computational methodologies. The flexible framework is planned to accommodate spatial transcriptomics, improved trajectory inference, and improved functionality for cell-cell communication analysis. Our open-source strategy invites contributions from the research community, with researchers able to craft bespoke modules while ensuring compatibility with the primary pipeline. This flexibility means that *cellSight* improves in tandem with methodological innovation in the field, allowing researchers to access state-of-the-art analysis without requiring extensive computational know-how.

## METHODS

Single-cell RNA sequencing (scRNA-seq) has transformed our ability to study cellular diversity by enabling the analysis of gene expression at the individual cell level^3,4^. However, the complexity of scRNA-seq data demands sophisticated analytical tools to derive meaningful insights^5,6^. To mitigate this challenge, *cellSight* integrates several key steps in single-cell genomics analysis. The process began with QC and normalization^19^, followed by dimension reduction and clustering to identify distinct cell populations^17^. We conducted differential expression analysis to identify gene expression changes between conditions^67^ and used cell-cell communication analysis to investigate how different cell types interact^79^. Canonical correlation analysis (CCA) was applied to integrate multiple datasets^22^, allowing for a comprehensive identification of clusters and biological states. *cellSight* also incorporates methodologies from *Seurat*^*80*^, enhancing its precision and versatility.

### DATA PREPROCESSING AND QC

scRNA-seq data undergoes stringent QC and preprocessing steps before analysis. Raw count matrices are loaded into *cellSight*, where standard QC procedures are implemented. These procedures include identifying and removing low-quality cells^19^ based on criteria such as mitochondrial content^20^, library size^81^, and unique molecular identifier (UMI) counts^21^.

Additionally, *cellSight* incorporates widely accepted QC metrics like mitochondrial percentage, number of detected features per cell, and total RNA molecule count per cell from the *Seurat*^*22*^ package, ensuring compatibility with established practices in single-cell analysis.

### NORMALIZATION

Normalization is crucial in ensuring the comparability of expression profiles across individual cells^82^. In *cellSight*, normalization is performed using the *sctransform* method^83^, which accounts for both technical noise and biological variability. In over-dispersed data, where gene expression variability exceeds the expectations of basic models like the Poisson distribution^84^, the increased variability stems from both biological differences between cells and technical factors, such as dropout events and batch effects^85,86^, making the data more challenging to model. In scRNA-seq, normalization typically involves adjusting the signal with a dropout parameter to account for these complexities^87^. This strategy has a major flaw of overfitting since different groups of genes are being normalized by the same factor^88^.

*Sctransform* uses a generalized linear model to estimate each gene UMI count and then pools similar genes with identical gene expression together to regularize the parameter estimate to produce error models^89^. This process of coupling various genes into groups and then estimating the error preserves the underlying biological dissimilarity by applying *sctransform*^*83*^, *cellSight* enhances the accuracy of downstream analyses, enabling robust identification of differentially expressed genes^90^.

### DIMENSION REDUCTION AND VISUALIZATION

Dimension reduction techniques are used to uncover the latent structure within single-cell data. Principal Component Analysis (PCA) is a key method integrated into *cellSight*, which maps the high-dimensional data to its principal components. The resulting components are utilized to visualize the data in a reduced-dimensional space, providing insights into the underlying cellular heterogeneity. A *k*-nearest neighbor (KNN) graph is then constructed based on similarities between cells in this reduced space; the edges in the graph represent connections between cells that share similar gene expression profiles. The Louvain algorithm^91^ is applied to this graph to identify clusters, representing groups of cells with similar expression patterns. These clusters can then be visualized and further analyzed to discover biological insights, such as identifying cell types or states. The resolution parameter controls the granularity of clustering, allowing users to define broader or finer groupings of cells.

### DIFFERENTIAL EXPRESSION ANALYSIS

Differential expression analysis is a cornerstone of single-cell studies, enabling the identification of genes that exhibit significant expression changes across different cell populations. In *cellSight*, this analysis is facilitated by the *Tweedieverse* statistical framework. *Tweedieverse* offers a flexible and robust tool for differential expression analysis, accommodating the unique characteristics of single-cell data and providing reliable results for the identification of genes driving cellular diversity. *Tweedieverse* uses the Tweedie model^92^ to discover differentially expressed genes. scRNA-seq is riddled with zero-inflated values, where the estimated coefficients are adjusted by considering the compound Poisson model, which is a part of the generalized Tweedie model.

While *DESeq2*^*93*^ has been widely adopted for differential expression analysis in RNA sequencing data, single-cell RNA sequencing (scRNA-seq) presents unique analytical challenges that necessitate more specialized statistical approaches ^13,94^. Our implementation of Tweedieverse in *cellSight* offers several key advantages over *DESeq2*, particularly in handling the characteristic zero-inflation of scRNA-seq data across the normalized interval values, where genes may show no expression in many cells due to both technical dropouts and true biological absence^87,95^. Unlike *DESeq2*’s negative binomial model, *Tweedieverse* implements a list of models, including the normal, Poisson, negative binomial, zero-inflated negative binomial, and compound Poisson distributions that naturally accommodate excess zeros while maintaining appropriate variance modeling, resulting in an improvement in sensitivity for detecting differentially expressed genes. Furthermore, *Tweedieverse*’s flexible parametric framework better captures cell-to-cell heterogeneity within defined populations [9], and its specialized normalization approaches are better suited for single-cell expression profiles compared to *DESeq2*’s size factor normalization^83,86^. Importantly, Tweedieverse’s computational efficiency scales better with increasing cell numbers, maintaining consistent performance across datasets ranging from 5,000 to 50,000 cells, while *DESeq2* shows substantial increases in computational overhead^96^, making it particularly well-suited for integration into *cellSight*’s automated workflow.

### CELL-CELL INTERACTION ANALYSIS

Understanding cellular communication is crucial for deciphering complex biological processes^97,98^. To address this, *cellSight* incorporates the *Cellchat*^*79*^ package, allowing for the analysis of cell-cell interactions based on ligand-receptor^99,100^ pairs. *cellSight* provides a comprehensive view of the communication network within a cellular population by integrating information on ligand-receptor interactions^101,102^.

### VISUALIZATION AND INTERPRETATION

The results obtained from the aforementioned analyses are visualized using various plotting techniques within the *cellSight* environment. These include t-distributed stochastic neighbor emulation (t-SNE) and uniform manifold approximation and projection (UMAP), which offer intuitive representations of cellular relationships and structures.

### GRAPH ATTENTION NETWORK ARCHITECTURE FOR SPATIAL ANALYSIS

The *cellSight* spatial transcriptomics module employs a novel Graph Attention Convolution (GATconv) framework to integrate gene expression profiles with spatial coordinate information, enabling comprehensive analysis of spatially-resolved transcriptomic data^103,104^. The methodology transforms traditional single-cell expression matrices into graph-structured data where individual cells or spots serve as nodes, each characterized by both transcriptional signatures and precise spatial positioning within the tissue architecture^105,106^.

The input data preprocessing pipeline converts single-cell expression profiles and corresponding spatial coordinates into a unified graph representation^107^. Each node in the constructed graph contains the complete gene expression vector for a given spatial location, while edges are established based on spatial proximity relationships determined through k-nearest neighbor algorithms^33,108^. This graph construction methodology preserves both the molecular characteristics of individual cells and their spatial context within the broader tissue organization^109^.

The core GATconv architecture implements self-attention mechanisms to dynamically weight the importance of neighboring nodes during feature aggregation^110,111^. The attention weights are computed through a learned transformation that considers both the gene expression similarity between neighboring cells and their spatial proximity^112,113^. This dual weighting system enables the model to identify spatially-constrained expression patterns while maintaining sensitivity to transcriptional relationships that may span larger spatial distances^114,115^.

### TRAINING AND OPTIMIZATION OF SPATIAL FEATURE LEARNING

The training procedure for the spatial module utilizes a multi-objective optimization framework that balances reconstruction accuracy with spatial coherence^116,117^. The model learns to predict gene expression patterns based on both local neighborhood information and broader spatial context through iterative attention-based message passing^118^. During each training iteration, nodes aggregate information from their spatial neighbors, with attention weights determining the relative contribution of each neighboring cell to the target node’s updated representation^2,119^.

The self-attention mechanism computes attention coefficients through a feedforward neural network that takes concatenated node features as input^120^. These coefficients undergo softmax normalization across all neighbors, ensuring that the attention weights sum to unity for each target node^121,122^. The resulting weighted feature aggregation enables the model to learn spatially-aware representations that capture both local microenvironmental effects and broader tissue-level organizational patterns^123,124^.

Performance evaluation during training employs multiple metrics including reconstruction accuracy, spatial clustering coherence, and biological pathway enrichment scores^125^. The optimization procedure incorporates spatial regularization terms that encourage similar expression patterns in spatially proximate regions while maintaining flexibility to detect sharp boundaries between distinct tissue domains or cell populations^126,127^.

### SPATIAL DOMAIN IDENTIFICATION AND CELLULAR COMMUNICATION NETWORKS

The trained GATconv model generates spatially-aware embeddings that enable automated identification of tissue domains with distinct molecular signatures^120,128^. These embeddings undergo dimension reduction and clustering procedures specifically designed to preserve spatial contiguity while identifying biologically meaningful cell populations^129^. The resulting spatial domains represent regions of similar transcriptional activity that correspond to anatomically distinct tissue structures or functional cellular neighborhoods^130,131^.

Cell-cell communication analysis within the spatial framework leverages the learned attention weights to identify communication hotspots and directional signaling pathway^132^. The attention mechanisms inherently capture relationships between spatially proximate cells, providing quantitative measures of intercellular influence that can be interpreted as communication strength^133^. This approach enables systematic mapping of ligand-receptor interactions within their native spatial context, revealing communication networks that would be undetectable through conventional single-cell analysis approaches^134,135^.

The spatial communication networks generated by the GATconv module identify both short-range paracrine signaling between immediate neighbors and longer-range communication patterns that span multiple cell diameters^136,137^. These multi-scale communication maps provide insights into tissue organization principles and reveal how cellular neighborhoods coordinate their molecular activities to maintain tissue homeostasis or respond to environmental perturbations^138,139^.

### INTEGRATION WITH EXISTING TOOLS

*cellSight* seamlessly integrates with established tools such as *Seurat*, ensuring compatibility with widely adopted workflows in the single-cell analysis community. This integration allows users to leverage the strengths of both *cellSight* and *Seurat*, expanding the analytical capabilities and providing a more comprehensive toolkit for researchers.

In summary, *cellSight* amalgamates state-of-the-art methods for QC, normalization, dimension reduction, differential expression analysis, and cell-cell interaction analysis, offering a robust and comprehensive solution for unraveling the complexities of single-cell genomics.

Finally, to assess the performance of *cellSight*, we conducted thorough evaluations on two independent skin-related studies. The first dataset, derived from mouse skin tissue 24 hrs after injury, focused on how the different cell type dynamics change to adapt and heal the wound. The second dataset, originating from human skin, centered around the importance of fibroblasts’ role in skin aging. Performance metrics, including QC plots, integration plots, and clustering, were calculated to validate the tool’s effectiveness in accurately capturing cellular heterogeneity and differential gene expression patterns within the skin-related contexts.

## RESOURCE AVAILABILITY

### LEAD CONTACT

Further information and requests for resources and analysis should be directed to and will be fulfilled by the lead contact, Ali Rahnavard.

### DATA AND CODE AVAILABILITY

All original code has been deposited at https://zenodo.org/records/10041147 and is publicly available at https://github.com/omicsEye/cellSight.

Any additional information required to reanalyze the data reported in this paper is available from the lead contact upon request.

## ACKNOWLEDGMENTS

The methodology development of this work was supported by the National Science Foundation grant DEB-2109688 to A.R. and application investigation as part of effort supported by an NIH grant to B.A.S. and A.R. from NIAMS (AR082417).

GEO accession GSE147744 and GSE265996.

## AUTHOR CONTRIBUTIONS

Conceptualization, A.R. and B.A.S.; methodology, A.R., R.C, and C.G.; investigation, A.R., C.G., A.R., and B.A.S.; writing, original draft, R.C., C.G., A.R.T., and A.R.; writing, review & editing, R.C., A.R., C.G., and B.A.S.; funding acquisition, B.A.S., and A.R.; supervision, A.R., A.R.T., and B.A.S.

## DECLARATION OF GENERATIVE AI AND AI-ASSISTED TECHNOLOGIES

The author(s) utilized ChatGPT and Claude to help refine sentence structure during the preparation of this work. Following the use of this tool, the author(s) thoroughly reviewed and edited the content as necessary, retaining full responsibility for the final content of the publication.

